# Salt sweetens the deal: bumble bees prefer sodium-enriched nectar when its sugar concentration is low

**DOI:** 10.64898/2026.09.19.752816

**Authors:** Amanda Vieira da Silva, Avery L. Russell

**Affiliations:** Centro de Ciências Naturais e Humanas, Universidade Federal do ABC, São Bernardo do Campo, São Paulo, Brazil; Department of Biology, Missouri State University, Springfield, MO, 65897, USA

**Keywords:** foraging, preference, micronutrients, sodium, floral rewards, decision making

## Abstract

Foraging decisions often depend on net energy intake, but the need for specific micronutrients means that foods are not always comparable based on energy content. Although sodium is essential for many physiological functions, many organisms face challenges obtaining it, especially those relying on a plant-based diet. Flowering plants often lure pollinators by manipulating nectar carbohydrate content, but nectar sodium might also influence pollinator choice. We experimentally tested how sodium addition and nectar carbohydrate concentration interact to affect bumble bee foraging decisions by allowing bees to forage on artificial flowers containing either low- or high-sucrose nectar, with or without sodium. We also explored whether sugar and sodium content in floral nectar trade-off across plant species. Bees preferred sodium-enriched nectar and preference nearly doubled when sucrose concentration was low. Furthermore, bees switched more often from flowers without sodium to those with sodium when sucrose concentration was low. Given increased fidelity for artificial flowers with nectar sodium, sodium may enhance plant fitness by increasing conspecific pollen receipt. Yet despite behavioral evidence that sodium enhances the perceived quality of low-sucrose nectar for bees, we found no evidence for a nectar sugar-sodium trade-off across plant species. We discuss implications for plant-pollinator interactions and flower evolution.

## INTRODUCTION

Animals require a variety of nutrients to achieve a nutritional balance that maximizes growth, survival and reproduction (Raubenheimer and Simpson 2018). However, achieving this balance is challenging because nutrient composition varies among food types (Raubenheimer and Simpson 2004, Filipiak et al. 2023). To overcome this challenge, animals often regulate how much of each food type they eat, including consuming micronutrient-rich food types that simultaneously contribute little or not at all to energetic intake (Simpson and Raubenheimer 1993). Sodium is one such essential micronutrient that is necessary for many physiological functions, including osmoregulation, muscle contraction and relaxation, conduction of nerve impulses, and DNA stability (Fraústo da Silva and Williams 2001, Kaspari 2021). Despite its physiological importance, obtaining sodium is difficult for many organisms, especially for those relying on a plant-based diet (Welti et al. 2019, Kaspari 2020, Demi et al. 2021). This limitation in acquiring sodium affects animal physiology, behavior, and performance (Snell-Rood et al. 2014, Raubenheimer and Simpson 2018). For example, to acquire sodium, herbivorous mammals are frequently observed licking salt from minerals and consuming soil (Duvall et al. 2023) and herbivorous arthropods are observed puddling (Molleman 2010, Demi et al. 2021), even though these nutrient sources do not contribute to dietary energy. Despite many studies showing that herbivorous animals can evaluate sodium content and exhibit sodium-seeking behaviors (Bonoan et al. 2017, Demi et al. 2021, De Sousa et al. 2022, Duvall et al. 2023), how sodium content affects choice of food types differing in nutritional content remains poorly understood (but see Clay et al. 2017, Borer et al. 2019)

In plant-pollinator mutualisms, flowering plants frequently offer carbohydrate-rich nectar to entice animals to visit and thereby acquire and transfer pollen (Nepi 2017, Ballarin et al. 2024). In contrast, the sodium content of floral nectar is very low, although highly variable across plant species, ranging from 0 to 35.7 mM (Hiebert and Calder 1983, Nicolson and W-Worswick 1990, Kaspari 2020, Filipiak et al. 2023). As a result of the low concentration of sodium in floral nectar, pollinators may suffer nutritional imbalance and sodium deprivation (Demi et al. 2021). For instance, honey bees allowed to self-select for the sodium concentration in their diet show a strong preference for high sodium concentrations, ranging from 21.7 to 43.5 mM (De Sousa et al. 2022), which are typically well above the concentrations found in most floral nectar (Hiebert and Calder 1983, Nicolson and W-Worswick 1990, Filipiak et al. 2023). As a result, pollinators are expected to employ foraging strategies to cope with sodium deprivation imposed by their plant diet. For example, pollinators might sample the floral nectar of multiple plant species and select those that meet their sodium intake requirements (Pyke 2010). In fact, recent field experiments have found that pollinators strongly prefer to forage on experimentally sodium-enriched nectar, resulting in an increase in the frequency of visits and the diversity of pollinators on sodium-enriched flowers (Finkelstein et al. 2022, Lovett and Carr 2024, VanValkenburg et al. 2024).

At the same time, the sugar content of nectar varies greatly including among flowers of a single plant (Herrera et al. 2006), across plant populations, and among species (Nicolson 2022, Liu et al. 2024). For instance, nectar sugar concentration is as low as 159.74 mM in *Lutheria splendens* (Bromeliaceae) and as high as 2707.80 mM in *Billbergia euphemiae* (Bromeliaceae) (Göttlinger and Lohaus 2022b). Although environmental factors (Brito Vera and Pérez 2024) and even floral microbes (Vannette 2020) can contribute to variation in nectar sugar concentration, the cost of floral nectar production may also play a role (Southwick 1984, Pyke 1991, Ornelas et al. 2007, — but see Pyke and Ren 2023). As a result, there is often strong selection for plants to manipulate their pollinators, such as by secreting colored or even pharmacologically active secondary metabolites into nectar (Nepi 2017, Parachnowitsch et al. 2019). Such manipulants can increase pollinator preference and consecutive conspecific visitations that can facilitate plant reproduction (i.e., floral fidelity) (Stevenson 2020). Could nectar sodium content also act as a manipulant of pollinator behavior? Accordingly, we might expect that pollinator preference and fidelity for a flower type may be enhanced when sodium content of its nectar increases, even when its sugar content decreases (Kaspari 2020). Yet pollinators often exhibit strong preferences for increased nectar sugar content (Nicolson et al. 2007, Fowler et al. 2016). An alternative possibility is thus that increased sodium content could enhance perceived nectar quality regardless of sugar content (Petanidou et al. 2006, Kaspari et al. 2008, Kaspari 2020).

In this laboratory study, we experimentally tested how nectar quality affected bumble bee (*Bombus impatiens* Cresson 1863) foraging decisions, by manipulating sucrose concentration and sodium presence in the nectar of artificial flowers. We hypothesized that if sodium is a floral nectar manipulant, sodium-enriched nectar will be preferred by bees over nectar lacking sodium, and this preference should be stronger when sugar content is low (Hypothesis 1; Figure 1). We therefore predicted that bumble bees will be more prone to (a) land and reject flowers with a low sugar content and no sodium when compared to flowers with a low sugar content and sodium; (b) reject nectar with low sugar content lacking sodium relative to nectar with high sugar content, regardless of sodium content, and (c) forage preferentially on sodium-enriched nectar and this preference will be stronger when bees are offered nectar with low sugar content. Accordingly, we hypothesized that bumble bees will be less prone to switch to a different flower type (i.e., exhibit greater floral fidelity) after tasting nectar with sodium, and switching will be reduced when bees are offered nectar with low sugar content (Hypothesis 2; Figure 1). Finally, we quantified the relationship between nectar sugar and sodium concentration across plant species using published data. We hypothesized that plant species that offered less sugar in their nectar would offer more sodium in their nectar (Hypothesis 3; Figure 1), as part of an evolved strategy to enhance the quality of their nectar and manipulate their pollinators.

**Figure 1.**
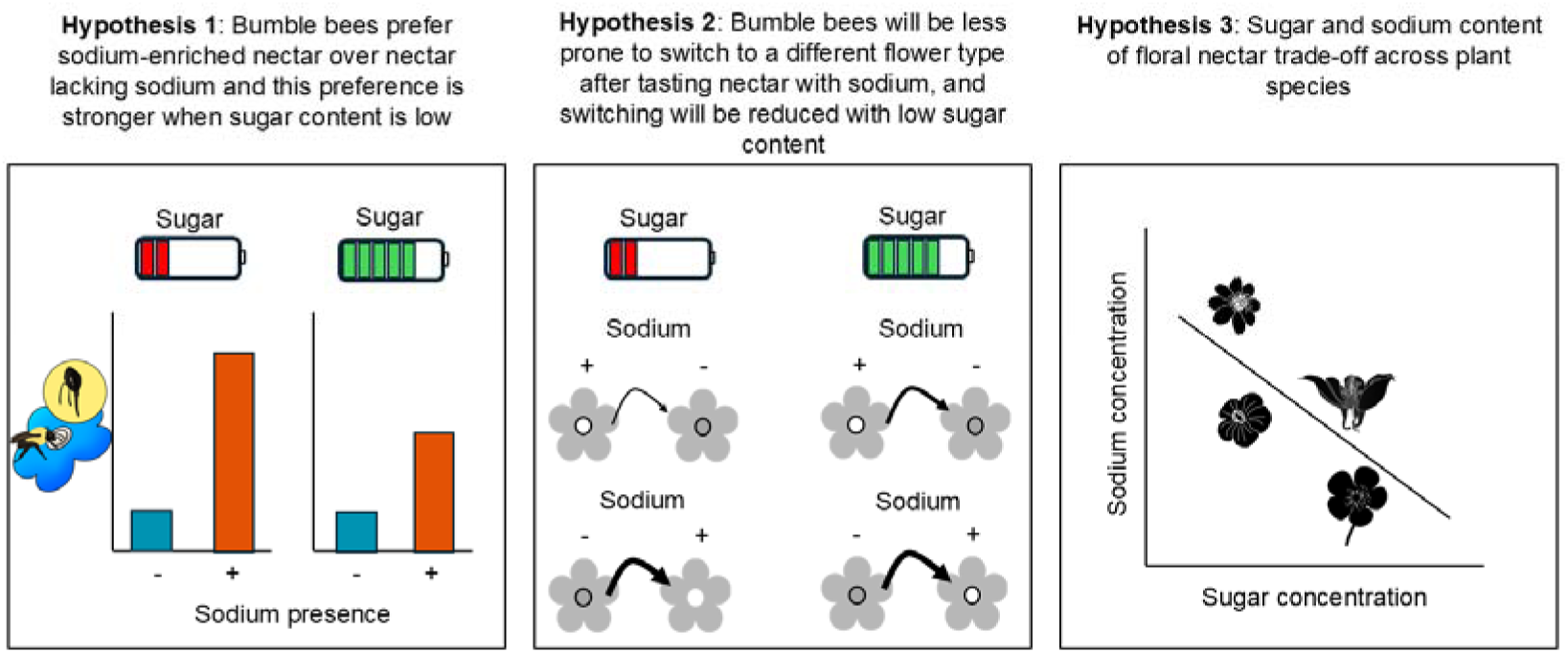
Schematic representation of the three proposed hypotheses. Hypothesis 1: sodium-enriched nectar will be preferred by bees over nectar lacking sodium, and this preference will be stronger when nectar sugar content is low. Hypothesis 2: bees will exhibit greater floral fidelity to flowers with sodium-enriched nectar, with floral fidelity being greatest when nectar sugar content is low. Hypothesis 3: plant species that offer less sugar in their nectar offer more sodium in their nectar.

## METHODS

### Test subjects

To evaluate how bumble bee foraging decisions are affected by nectar quality and sodium addition, we maintained two colonies (Plant Products: Biobest Group, Canton, MI, U.S.A.) of the common eastern bumble bee *Bombus impatiens* following Russell et al. (2017), and Russell et al. (2020). In brief, we allowed colonies to forage freely on 20% w/v sucrose solution from artificial feeders within enclosed foraging arenas (length, width, height: 82 × 60 × 60 cm) set to a 14 h : 10 h light : dark cycle. Each foraging arena contained two artificial feeders. Each feeder’s plastic lid was half blue and half yellow to familiarize foraging bees with these colors. Pulverized honeybee-collected pollen (Koppert Biological Systems) was deposited within colony boxes (2g / day).

### Experiment

To identify appropriate test subjects and accustom bees to foraging for nectar on our experimental set, we captured bees that foraged on the artificial feeders and marked them with non-toxic oil markers (Sharpie, CA) and returned them to their colonies. We considered that they were appropriate for experiments when marked bees were observed foraging again on the artificial feeders.

To assess how sodium and sucrose concentration of nectar affected bee foraging behavior, we divided 36 marked flower-naïve bees into two treatment groups, each with two sub-treatments. Both colonies were equally represented per treatment and sub-treatment. Across all trials we set up a 5 × 4 horizontal array of equal numbers of cleaned (wiped with 70% ethanol between trials and air dried) yellow and blue colored plastic flowers on the test arena floor (20 flowers total). Each flower received an artificial nectary (a sterile 1.5 mL microcentrifuge cap), into which 3 µL of sterile artificial nectar solution was added into the center of the cap. The two treatment groups differed in terms of which sucrose concentration was present (20% or 50% w/v; 584.3 mM or 1460.7 mM, respectively; Figure 2). The sub-treatments differed in terms of which one of the two flower colors (i.e., flower types) offered nectar with 0.1% NaCl w/v (17.11 mM) (Fisher Scientific) (Figure 2). We chose these ecologically realistic sucrose and salt concentrations based on the range of sucrose and NaCl concentrations found in the nectar of live plants (sugar concentration reported in nectar of live flowers ranges from 159.74 mM to 2707.80 mM (Göttlinger and Lohaus 2022b); while sodium concentration ranges from 0 to 35.7 mM; Hiebert and Calder 1983, Nicolson and W-Worswick 1990, Kaspari 2020, Filipiak et al. 2023; see results). The flower color type that offered NaCl was systematically alternated between behavioral trials. We used two flower colors in each horizontal array of 20 flowers to facilitate bees being able to choose between the nectar solution types (i.e., by associating a given flower color with NaCl or no NaCl; bumble bees rapidly learn simple flower color-nectar reward associations - Guiraud et al. 2025). We used a water purification system (Millipore, Sigma Aldrich) to prepare deionized purified sterile water for solutions.

**Figure 2.**
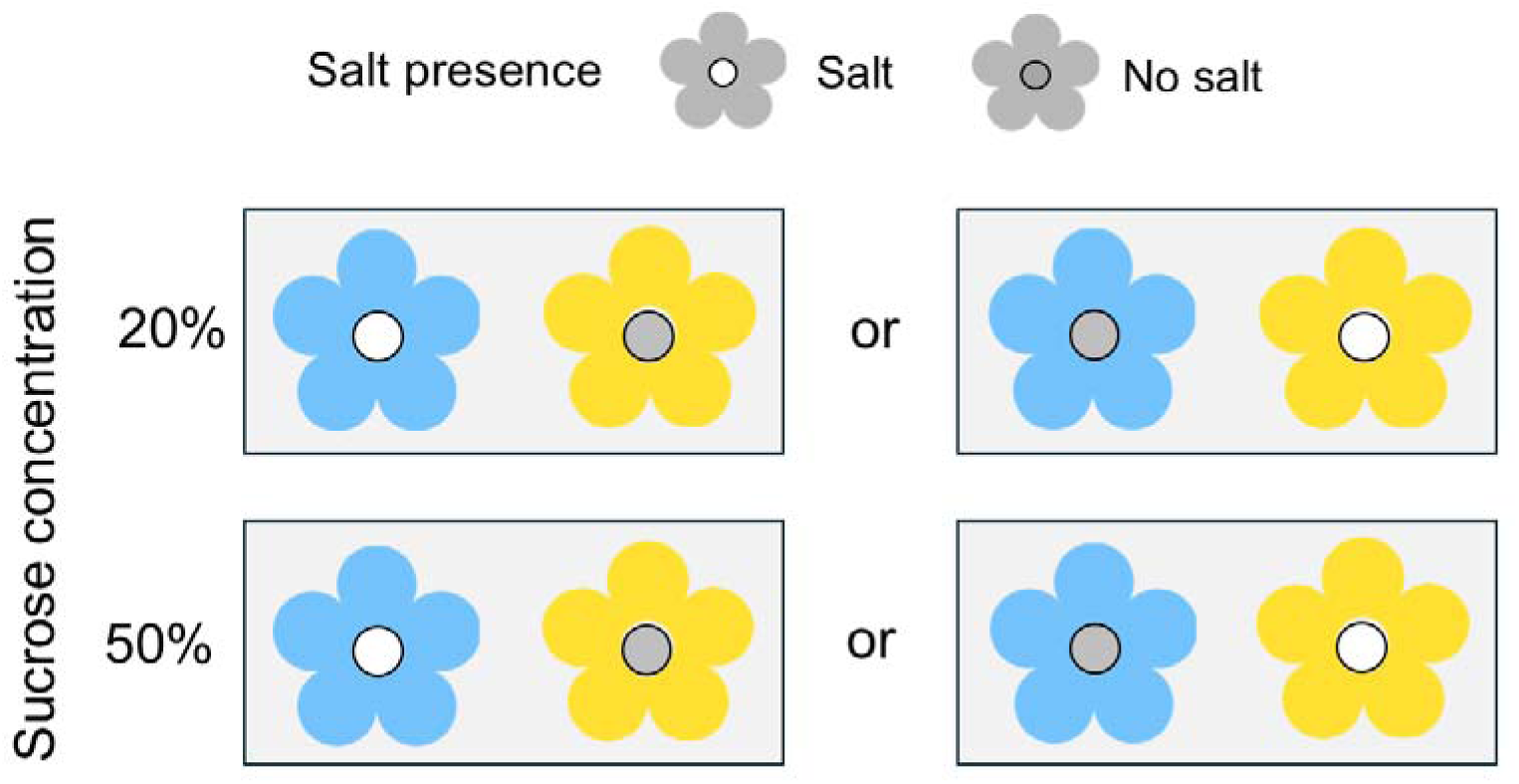
Experimental set up. Each bumble bee (*N* = 36) was individually tested using an array of 20 artificial flowers with either all low-sugar (20% w/v sucrose treatment; 584.3 mM) or all high-sugar nectar (50% w/v sucrose treatment; 1460.7 mM). Within each treatment, bees had access to equal numbers of both flower color types alternating by position in the array, one type with 0.1% w/v NaCl salt (17.11 mM) and the other type with no NaCl in their nectar. The color of the flower (blue or yellow) associated with NaCl differs among sub-treatments.

To initiate a behavioral trial, freshly cleaned flowers and nectaries were set up and a single marked worker bee was gently captured from the foraging arena using a 40 dram vial (Bioquip, CA) and immediately released in the center of the test arena following Russell et al. (2017). We recorded three flower visiting behaviors: ‘landing’, when the bee visited the flower (touched the flower with at least 3 of its legs simultaneously), but did not extend its proboscis into the nectary; ‘tasting’, when the bee visited and extended its proboscis into the nectary only briefly (<<1 second; leaving most of the nectar); and ‘drinking’, when the bee visited and extended its proboscis into the nectary for more than 1 second (removing all of the nectar).

Immediately after each visit that involved drinking, we refilled the artificial nectary with the appropriate nectar solution, such that flowers were never depleted within a trial. Bees made on average 29.6 ± 13.5 flower visits (mean ± standard deviation) within their single trial. To ensure trials were comparable, we terminated a trial when the bee did not approach any flower for a period of 5 min. To terminate a trial, we turned off the overhead arena lights and captured the bee in a vial. After a bee completed its trial, it was euthanized to prevent it from returning experimental nectar to its colony. After each trial, flowers were wiped clean with 70% ethanol and artificial nectaries were soaked in 70% ethanol for at least 30 minutes and then rinsed repeatedly in sterile water and allowed to air dry before being reused.

### Literature review

To evaluate whether plants increase the amount of sodium available in their nectar to improve the quality of a nectar with low sugar content, we searched for papers quantifying the amount of sodium and sugar in the nectar of live flowers under field and cultivated conditions. For that, we conducted a systematic search on Web of Science (main core collection) for published papers between 1945 (first year available on Web of Science) and May 14^th^, 2026. We used the following keywords: (1) salt AND floral nectar; (2) sodium AND floral nectar; (3) NaCl AND floral nectar; (4) (sodium OR salt OR NaCl or ion) AND floral nectar composition; and (5) floral nectar AND (sodium OR salt OR NaCl or ion). We also included in our database all studies that cited Hiebert and Calder (1983) and Nicolson and W-Worswick (1990), because these studies represent the foundation of understanding sodium availability on flowers. We also included studies that cited Finkelstein et al. (2022), since this is one of the first studies to manipulate sodium content of floral nectar in a natural community. Finally, to be as thorough as possible, we searched through the references of these studies. Therefore, our final database comprised 149 studies. We read these studies and included in our dataset only studies that provided data on sodium and sugar content on floral nectar. For studies that manipulated sodium or sugar content, we only included those that provided data for the control conditions (no sodium or sugar manipulation). For each study, we collected plant species, sugar and sodium content. We synonymized plant species names using Plants of the World Online (POWO 2024).

### Statistical analysis

All data were analyzed using R v. 4.1.3 (R Core Team 2022). We checked model assumptions using *DHARMa* v. 0.4.6 package (Hartig 2020) and conducted visual diagnostics with *sjPlot* (Lüdecke 2023). For figures, we used *ggplot2* v. 3.5.1 (Wickham 2016).

To evaluate whether sodium and sugar concentration affected bee foraging preference, we specified three generalized linear mixed effects models (GLMMs). As the response variable, we used the (1) proportion of landings (for a given trial: total visits with landing / total visits with landing plus total visits with drinking), (2) proportion of tasting (for a given trial: total visits with tasting / total flower visits with drinking plus total visits with tasting), or (3) the proportion of drinking (for a given trial: total visits with drinking / total flower visits). Each response variable was used in one GLMM. To evaluate whether bees switched to a different flower type while foraging, we conducted a GLMM using the proportion of switching to a different flower type compared to the immediate previous visit as the response variable. We used the interaction between sucrose concentration and the presence of salt as the predictor variable in all models. We included the identity of the bee as a random variable in all models.

For all models, we considered a binomial family distribution using the *glmmTMB* function available in *glmmTMB* v. 1.1.7 package (Brooks et al. 2017). To generate p-values, we specified type II Wald χ²-tests via the *Anova* function available in *car* v. 3.1-2 package (Fox and Weisberg 2019). If we found an effect of the predictor variable, we used an adjusted Tukey’s post-hoc test to evaluate the difference between the levels of the predictor using the *emmeans* function available in the *emmeans* v. 1.8.8 package (Lenth et al. 2025). We calculated effect sizes using the *emmeans* package (Lenth 2025) and reported Cohen’s d in all cases.

To evaluate the relationship between sodium and sugar concentration in the nectar of live plants, we conducted a phylogenetic generalized least square (PGLS) model to account for evolutionary relationships among plant species. We used the mean values of sodium (mM) and sugar concentration (mM) for each species. We used the largest dated angiosperm phylogenetic tree at the species level by Janssens et al. (2020). This tree was reconstructed using 36,101 angiosperm species based on two plastid barcoding genes, matK and rbcL. It was dated using 56 angiosperm fossils as calibration points. Before running the analysis, we pruned all phylogenetic trees to species whose sodium and sugar concentrations were simultaneously quantified. We used this approach because nectar concentration is highly variable not only across individual plants populations, but also across individual flowers (Parachnowitsch et al. 2019). To create our regression model, we used the *phylolm* in *phylolm* v. 2.6.2 (Tung Ho and Ané 2014). To account for phylogenetic relatedness, we incorporated a correlation structure in our regression. This structure assumes that the variance of trait values increases linearly with time. To conduct all analysis, we also used *phytools* v. 1.9-16 (Revell 2012), *ape* v. 5.7-1 (Paradis and Schliep 2019), *geiger* v. 2.0.11 (Pennell et al. 2014) and *caper* v. 1.0.1 (Orme et al. 2018).

## RESULTS

### Bumble bees prefer sodium-enriched flowers

Bumble bees rejected artificial flowers (landing on them without drinking or tasting) that lacked salted nectar much more frequently than flowers with salt (0.1% NaCl) nectar. This pattern only became very strong when the sucrose concentration of nectar was low (20% vs 50%) (Figure 3a; GLMM: sugar effect: χ1² = 4.735, *P* = 0.029; salt presence effect: χ1² = 12.494, *P* < 0.0001; sugar × salt presence effect: χ1² = 3.184, *P* = 0.074; effect size_low sugar_ = 0.876; effect size_high sugar_ = 0.289; Table S1). In fact, bees were 2.08 times as likely to reject flowers with low sucrose concentration lacking salt versus those with salt. Likewise, bees rejected nectar lacking salt after tasting much more frequently when the sucrose concentration of that nectar was low (Figure 3b; GLMM: sugar effect: χ1² = 20.269, *P* < 0.0001; sodium presence effect: χ1² = 34.347, *P* < 0.0001; sugar × salt presence effect: χ1² = 18.139, *P* < 0.0001; effect size_low sugar_ = 2.323; effect size_high sugar_ = −0.269; Table S1). Overall, bees were 5.93 times as likely to reject nectar lacking salt versus nectar with salt when sucrose concentration was low. Altogether, bees preferred to drink from flowers with salt, but this effect became much greater (1.73 times as likely) when sucrose concentration of the nectar was low (Figure 3c; GLMM: sugar effect: χ1² = 18.860, *P* < 0.0001; salt presence effect: χ1² = 40.855, *P* < 0.0001; sugar × salt presence effect: χ1² = 19.802, *P* < 0.0001; effect size_low sugar_ = −1.574; effect size_high sugar_ = −0.259; Table S1).

**Figure 3.**
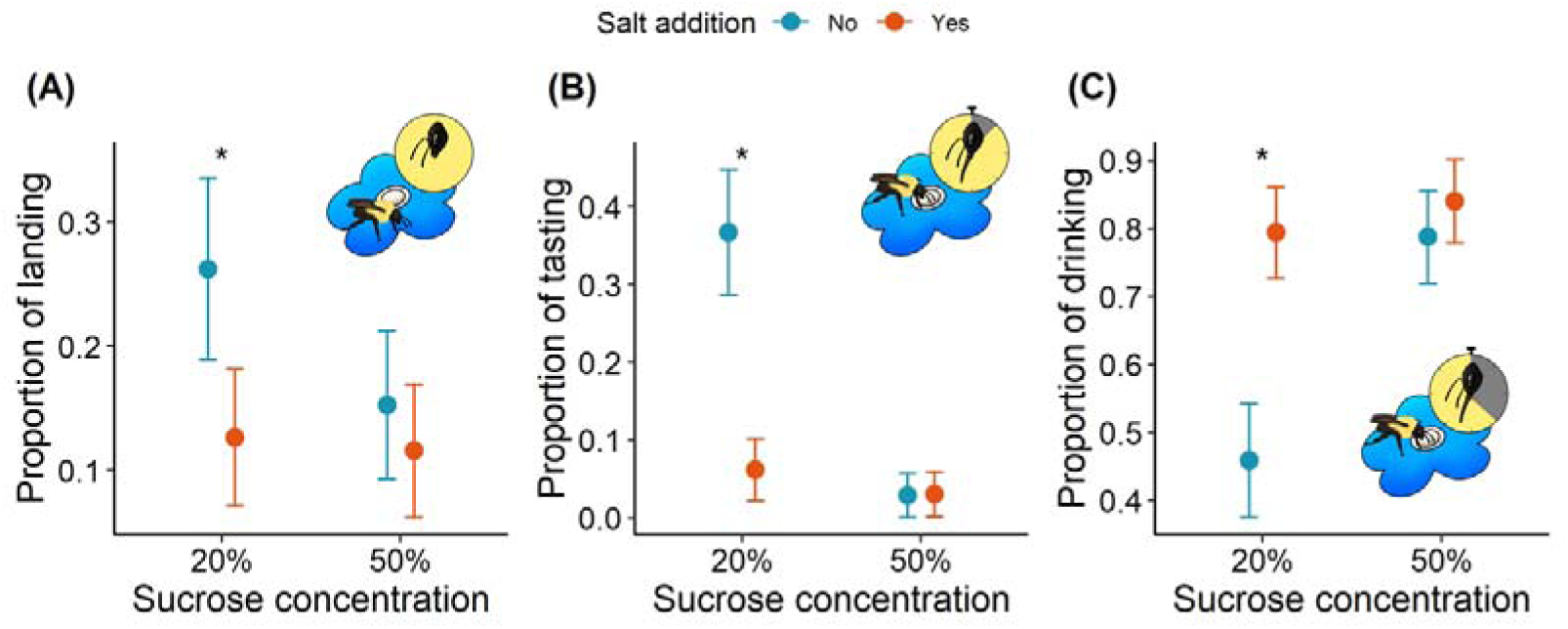
Effect of sucrose concentration (20% or 50% w/v; 584.3 mM or 1460.7 mM, respectively) and NaCl (0.1% w/v; 17.11 mM) presence on the proportion of (A) landings (visits without drinking or tasting), (B) tasting (visits without drinking), and (C) drinking by bumble bees (*Bombus impatiens*). A ‘landing’ was when the bee visited the flower (touched the flower with at least 3 of its legs simultaneously) but did not extend its proboscis into the nectary; ‘tasting’, when the bee visited and extended its proboscis into the nectary only briefly (<<1 second); and ‘drinking’, when the bee visited and extended its proboscis into the nectary for more than 1 second, removing all of the nectar. *N* = 18 bees for each treatment. Dots represent the mean values and lines represent the standard errors. Asterisks indicate significant differences in mean proportions among flower types for a given treatment at p < 0.05 according to a Tukey’s post hoc test. Illustrations by Matheus Siqueira.

The probability of switching to a different flower type was affected by sucrose concentration and salt presence (Figure 4; GLMM: sugar effect: χ1² = 0.505, *P* = 0.477; salt presence effect: χ1² = 0.018, *P* = 0.89; sugar × salt presence effect: χ1² = 4.848, *P* = 0.003; effect size_low sugar_ = 0.340; effect size_high sugar_ = 0.317; Table S1). Bees were 1.30 times more prone to switch from a flower lacking salt to a flower with salt when the sucrose concentration was low.

**Figure 4.**
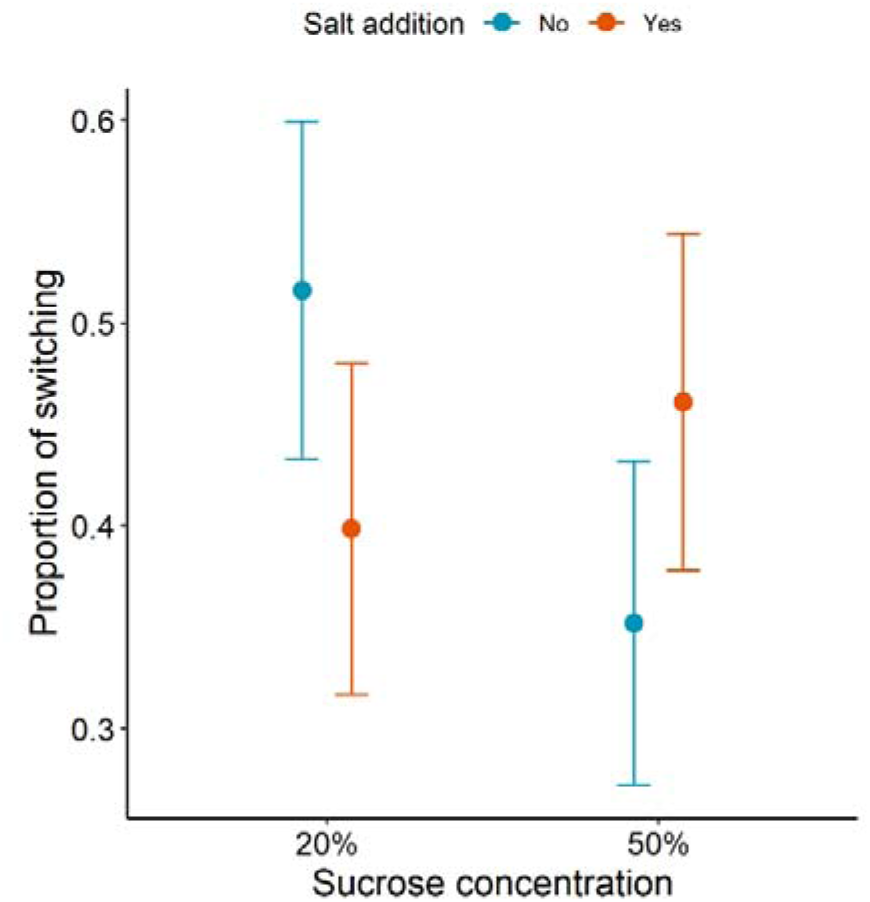
Effect of sucrose concentration (20% or 50% w/v; 584.3 mM or 1460.7 mM, respectively) and NaCl (0.1% w/v; 17.11 mM) presence on the proportion of switching to a different flower type (one with versus without NaCl in its nectar) by bumble bees (*Bombus impatiens*). *N* = 18 bees for each treatment. Dots represent the mean values and lines represent the standard errors.

### No phylogenetic relationship between sugar and sodium content

Most studies did not provide data on floral nectar composition. Only 15 of the 149 studies recovered in our literature search provided data on nectar sodium and/or sugar content available in live flowers. Of these, only eight studies provided data on both sodium and sugar content. These studies collectively reported sodium and sugar concentration of nectar for 215 plant species across 40 genera and nine families. The most common species studied were *Pitcairnia recurvata* (Bromeliaceae) (*k* = 7 observations), followed by *P. corallina, P. longissimiflora, P. maidifolia* and *P. wendlandii* (*k* = 6 observations for each species). The most common genera studied were *Pitcairnia* (*k* = 121 observations), *Aechmea* (Bromeliaceae) (*k* = 37), *Tillandsia* (Bromeliaceae) (*k* = 36). The most commonly studied family was Bromeliaceae (*k* = 347 observations) and Solanaceae (*k* = 22 observations). Unfortunately, two studies reported concentration of sugar using g/100 mL, but did not specify which sugar type was available in the sample (Nicolson 1990, Nicolson and W-Worswick 1990). Since we need sugar composition to convert this data to mM, we removed these studies from our analysis.

The mean sodium concentration found in the nectar of live flowers was 1.70 ± 4.44 mM (mean ± standard deviation) (median = 0.26 mM), whereas mean sugar concentration was 939 ± 432 mM (median = 894 mM). However, in our phylogenetic analysis we only included studies that measured the concentration of sodium and sugar in the same flower. Thus for studies included in our phylogenetic analysis, the mean sodium concentration was 0.61 ± 1.13 mM (median = 0.21 mM) while the mean sugar concentration of nectar was 935 ± 429 mM (median = 892 mM). To account for the phylogenetic relatedness among species, we pruned the angiosperm tree to our data (Janssens et al. 2020). Only 73 of the 215 species with data on sodium and sugar concentration of floral nectar were present on the tree. We found no significant relationship between sodium and sugar concentration in nectar across plant species (Figure 5A; PGLS with Brownian motion correlation structure: intercept = 0.14, *P* = 0.70; slope = 0.00012, *P* = 0.38). Accounting for nectar volume, flower nectar had on average 7.65 ± 21.6 ng of sodium (*N* = 8 species).

**Figure 5.**
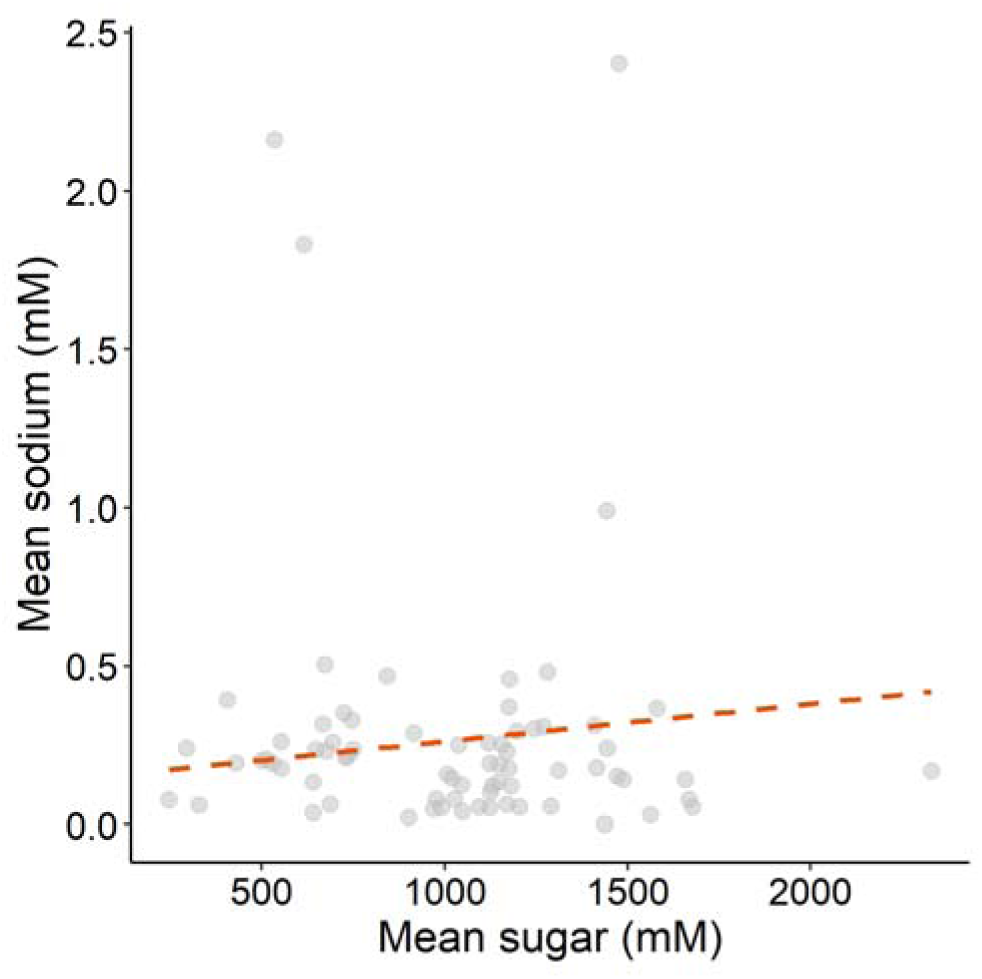
Correlated evolution between mean sugar and sodium concentration (mM) on live flowers of 73 plant species. Each dot represents the mean value of a plant species. Darker dots indicate overlapping data points. The orange dashed line represents the regression line estimated from the phylogenetic generalized least squares model in which the p-value of the slope is greater than 0.05.

## DISCUSSION

Because pollinators are expected to evaluate the nutrient qualities of their foods to achieve nutritional balance (Welti et al. 2019, Kaspari 2020, Demi et al. 2021), we hypothesized that access to a rare nutrient (i.e., sodium) would act as a manipulant of pollinator behavior and thus affect the perceived quality of nectar differing in energetic content (sucrose, a common nutrient). We found that bumble bees prefer sodium-enriched nectar of artificial flowers, consistent with recent studies conducted under field (Finkelstein et al. 2022, VanValkenburg et al. 2024) and experimental conditions (De Sousa et al. 2022). Furthermore, as we hypothesized, we found that the magnitude of sodium preference depended on sugar content (consistent with Hypothesis 1; Figure 1). In addition to affecting bee preference, sodium-enrichment also affected bee movement among artificial flower types (i.e., floral fidelity), partly consistent with Hypothesis 2 (Figure 1). When flowers offered only nectar with low sugar content, bees switched more often when encountering flowers without sodium (i.e., exhibited greater floral fidelity to flowers with sodium). Conversely, when flowers offered only nectar with high sugar content, bees switched more often when encountering sodium enriched nectar, suggesting that higher energetic content of the nectar reward may result in sodium presence being perceived as mildly aversive. Finally, despite the clear effect of sodium enhancing the perceived quality of nectar with low sugar content, we found no evidence for a negative relationship between nectar sugar and sodium content across taxonomically diverse plant species (inconsistent with Hypothesis 3; Figure 1). Thus, plants likely do not reduce the sugar provision of their nectar while simultaneously increasing its sodium content as a strategy to enhance the quality of nectar and thereby manipulate their pollinators.

Overall, our results strengthen the case for sodium to be considered a floral reward offered in exchange for pollination services (see Kaspari 2020), adding sodium to a growing list of non-energetic nutrients that act as floral reward (Lau and Nieh 2016, Bonoan et al. 2017). To be considered a floral reward, the floral component must be used by animals and ensure repeated visitation to flowers (Simpson and Neff 1981). We found that bees preferred sodium-enriched flowers, which also required bees to discriminate which flower color was associated with sodium-enriched nectar. Our results thus suggest that bees may have learned to associate sodium-enriched nectar with flower color, although future work will be required to categorically demonstrate associative learning in this context. Likewise, Finkelstein et al. (2022) and VanValkenburg et al. (2024) showed that sodium-enriched flowers increased pollinator visitation at the community level. Our study extends their foundational work by showing that in addition to consumption, bees frequently rejected (i.e., tasted but did not consume) low-sucrose nectar when it lacked sodium; however, when sucrose concentration was high, nectar rejection was very rare regardless of sodium content. A similar result was found when phenylalanine, an essential amino acid not produced by bees, was added to nectar with low sugar content (Hendriksma et al. 2014). Taken together, components of nectar that contribute to nutritional requirements, but not energetic content, appear to manipulate pollinators into foraging for nectar with low energetic content.

Preference and learning can be powerful mechanisms driving selection on floral traits (Schiestl and Johnson 2013) and even speciation (Takimoto et al. 2022). Given that bees preferred sodium-enriched nectar and may even have learned to associate this nutrient with flower color, nectar sodium content and associated floral traits might be a target for pollinator-mediated selection (Parachnowitsch et al. 2019, but see Abrahamczyk et al. 2017). For instance, plants offering nectar with low sugar content in particular may experience strong pollinator-mediated selection for honest signals of sodium content. Such signals would facilitate pollinators in reliably visiting flowers with sodium-enriched nectar, similar to honest signals associated with other kinds of floral rewards (Knauer and Schiestl 2015, Ortiz et al. 2020). Furthermore, while sodium content of nectar is on average low, it is typically much higher in pollen (Nicolson, 1990). Given that bees can assess pollen lipid and protein content (Ruedenauer et al. 2016, Vaudo et al. 2016) and sodium content of floral nectar (De Sousa et al. 2022), it is plausible to assume that bees can also assess pollen sodium content. We might therefore expect pollinator-mediated selection on pollen sodium content and associated floral traits to be even stronger than within the context of nectar foraging.

Pollinator foraging decisions are often driven by nutritional demands, which could include visiting multiple plant species to acquire sufficient nutrients, resulting in greater interspecific pollen transfer and reduced plant fitness (Morales and Traveset 2008). We should therefore expect plants to manipulate reward composition to promote floral fidelity (Huang et al. 2015, Pyke 2016, Nepi et al. 2018). Consistent with this hypothesis, we observed that fidelity to a given flower type was enhanced by adding sodium to a low-sugar floral nectar. Why then did we not observe a negative evolutionary relationship between sodium and sugar content in floral nectar across plant species? There are at least two non-mutually exclusive mechanisms explanations. First, although it is often assumed that the production and secretion of carbohydrate-rich nectar is costly for the plant, this cost may be highly variable depending on the ecological context (Pyke and Ren 2023). When photosynthetic metabolism is constrained, such as water, light, or nutrient deficiency, carbohydrate production – and thus nectar production – also becomes limited (Pacini and Nepi 2007). Conversely, when photosynthetic metabolism is not constrained, nectar cost might be low or even negligible (Pyke and Ren 2023). Understanding whether the cost of carbohydrate production influences nectar sodium content will require studies evaluating flower nectar sugar and sodium content for plant populations or species varying in constraints of photosynthetic metabolism. Second, while pollinators are usually sodium-deprived, so are herbivores (Kaspari 2020). While additional sodium in nectar attracts pollinators (Finkelstein et al. 2022, VanValkenburg et al. 2024), potentially facilitating plant fitness, additional sodium may also attract florivores that decrease plant fitness (Snell-Rood et al. 2014). Extrapolating from studies demonstrating that increased leaf sodium content increases folivory (Prather et al. 2018, Welti and Kaspari 2021), it is plausible that increased nectar sodium content also increases florivory. Thus, while pollinators may exert selective pressure for increased sodium in floral nectar, herbivory may exert a selective pressure in the opposite direction. How plants balance using sodium to lure pollinators while avoiding attraction of herbivores remains an open question.

Although biological mechanisms may explain the absence of a relationship between sodium and sugar investment in nectar, the absence of this relationship may also be due to limited data. While more than 200,000 plant species produce nectar as a floral reward (Ballarin et al. 2024), we were able to compile data on sodium and sugar content for only 215 species and only a third were represented in the currently most comprehensive phylogenetic tree at the species level (Janssens et al. 2020). Although floral nectar is one of the primary floral rewards, floral nectar composition beyond sugars is still relatively understudied (Nicolson and Thornburg 2007, Barberis et al. 2023).

Altogether, our study suggests that sodium-enriched nectar plays an important, but context-dependent role in plant-pollinator interactions. Additionally, given anthropogenic increases in soil salinity, such as via road salt and groundwater infiltration (Findlay and Kelly 2011, Ivushkin et al. 2019), understanding consequences for floral rewards, pollinator foraging, and animal-mediated pollination will become ever more important. Experiments manipulating soil salinity have shown that when soil salinity is high, nectar sodium content of at least some plant species increases, in turn attracting more pollinators (Lovett and Carr 2024). At the same time, our results suggest that enhanced pollinator preference may not always increase pollination success, given that when sugar content of nectar was high, fidelity to sodium-enriched flowers actually decreased. Furthermore, even if sodium-enrichment generally enhances pollination (Snell-Rood et al. 2014), plants not adapted to salty soil conditions might overall have reduced fitness due to physiological stress (Equiza et al. 2017). Finally, although anthropogenic increases in soil salinity are common, the soil in some regions is naturally salty, especially in coastal areas (Jaiswal et al. 2022). Future phylogenetic and experimental studies will be required to understand how variation in soil salinity affects plant-pollinator interactions and potential tradeoffs among floral reward nutrient types.

## ETHICS APPROVAL

All bumble bee experimentation was carried out in accordance with the legal and ethical standards of the USA.

*Studies that provided data for sodium and/or sugar concentration: (Hiebert and Calder 1983, Nicolson 1990, Nicolson and W-Worswick 1990, Bradshaw and Bradshaw 1999, Varassin et al. 2001, Afik et al. 2006, Afik et al. 2014, Tiedge and Lohaus 2017, Göttlinger and Lohaus 2020, Göttlinger and Lohaus 2022a, Lovett and Carr 2024)

## Supporting information

Analyses and datasets

## REFERENCES

Afik, O., A. Dag, Z. Kerem, and S. Shafir. 2006. Analyses of avocado (*Persea americana*) nectar properties and their perception by honey bees (*Apis mellifera*). Journal of Chemical Ecology 32:1949–1963.

Afik, O., K. S. Delaplane, S. Shafir, H. Moo-Valle, and J. J. G. Quezada-Euán. 2014. Nectar minerals as regulators of flower visitation in stingless bees and nectar hoarding wasps. Journal of Chemical Ecology 40:476–483.

Ballarin, C. S., F. E. Fontúrbel, A. R. Rech, P. E. Oliveira, G. A. Goés, D. S. Polizello, P. H. Oliveira, L. Hachuy-Filho, and F. W. Amorim. 2024. How many animal-pollinated angiosperms are nectar-producing? New Phytologist 243:2008–2020.

Barberis, M., D. Calabrese, M. Galloni, and M. Nepi. 2023. Secondary metabolites in nectar-mediated plant-pollinator relationships. Plants 12:550.

Bonoan, R. E., T. M. Tai, M. Tagle Rodriguez, L. Feller, S. R. Daddario, R. A. Czaja, L. D. O’Connor, G. Burruss, and P. T. Starks. 2017. Seasonality of salt foraging in honey bees (*Apis mellifera*). Ecological Entomology 42:195–201.

Borer, E. T., E. M. Lind, J. Firn, E. W. Seabloom, T. M. Anderson, E. S. Bakker, L. Biederman, K. J. La Pierre, A. S. MacDougall, J. L. Moore, A. C. Risch, M. Schutz, and C. J. Stevens. 2019. More salt, please: global patterns, responses and impacts of foliar sodium in grasslands. Ecology Letters 22:1136–1144.

Bradshaw, S. D., and F. J. Bradshaw. 1999. Field energetics and the estimation of pollen and nectar intake in the marsupial honey possum, Tarsipes rostratus , in heathland habitats of South-Western Australia. Journal of Comparative Physiology B: Biochemical, Systemic, and Environmental Physiology 169:569–580.

Brito Vera, G. A., and F. Pérez. 2024. Floral nectar (FN): drivers of variability, causes, and consequences. Brazilian Journal of Botany 47:473–483.

Brooks, M. E., K. Kristensen, K. J. V. Benthem, A. Magnusson, C. W. Berg, A. Nielsen, H. J. Skaug, M. Mächler, and B. M. Bolker. 2017. glmmTMB balances speed and flexibility among packages for zero-inflated generalized linear mixed modeling. The R Journal 9:378.

Clay, N. A., R. J. Lehrter, and M. Kaspari. 2017. Towards a geography of omnivory: Omnivores increase carnivory when sodium is limiting. Journal of Animal Ecology 86:1523–1531.

De Sousa, R. T., R. Darnell, and G. A. Wright. 2022. Behavioural regulation of mineral salt intake in honeybees: a self-selection approach. Philosophical Transactions of the Royal Society B: Biological Sciences 377.

Demi, L. M., B. W. Taylor, B. J. Reading, M. G. Tordoff, and R. R. Dunn. 2021. Understanding the evolution of nutritive taste in animals: Insights from biological stoichiometry and nutritional geometry. Ecology and Evolution 11:8441–8455.

Duvall, E. S., B. M. Griffiths, M. Clauss, and A. J. Abraham. 2023. Allometry of sodium requirements and mineral lick use among herbivorous mammals. Oikos 2023:e10058.

Equiza, M. A., M. Calvo-Polanco, D. Cirelli, J. Señorans, M. Wartenbe, C. Saunders, and J. J. Zwiazek. 2017. Long-term impact of road salt (NaCl) on soil and urban trees in Edmonton, Canada. Urban Forestry & Urban Greening 21:16–28.

Filipiak, Z. M., J. Ollerton, and M. Filipiak. 2023. Uncovering the significance of the ratio of food K:Na in bee ecology and evolution. Ecology 104.

Findlay, S. E. G., and V. R. Kelly. 2011. Emerging indirect and long-term road salt effects on ecosystems. Ann N Y Acad Sci 1223:58–68.

Finkelstein, C. J., P. J. Caradonna, A. Gruver, A. Ellen, M. Kaspari, and N. J. Sanders. 2022. Sodium-enriched floral nectar increases pollinator visitation rate and diversity. Biology Letters 18.

Fowler, R. E., E. L. Rotheray, and D. Goulson. 2016. Floral abundance and resource quality influence pollinator choice. Insect Conservation and Diversity 9:481–494.

Fox, J., and S. Weisberg. 2019. An R compation to applied regression. Sage, Thousand Oaks, California, USA.

Fraústo da Silva, J., and R. Williams. 2001. Sodium, potassium, and chlorine: osmotic control, electrolytic equilibria, and currents. Pages 231–249 in O. U. Press, editor. The biological chemistry of the elements: the inorganic chemistry of life. Oxford University Press, New York.

Göttlinger, T., and G. Lohaus. 2020. Influence of light, dark, temperature and drought on metabolite and ion composition in nectar and nectaries of an epiphytic bromeliad species (*Aechmea fasciata*). Plant Biol (Stuttg) 22:781–793.

Göttlinger, T., and G. Lohaus. 2022a. Comparative analyses of the metabolite and ion concentrations in nectar, nectaries, and leaves of 36 bromeliads with different photosynthesis and pollinator types. Frontiers in Plant Science 13.

Göttlinger, T., and G. Lohaus. 2022b. Comparative analyses of the metabolite and ion concentrations in nectar, nectaries, and leaves of 36 bromeliads with different photosynthesis and pollinator types. Frontiers in Plant Science 13:987145.

Guiraud, M.-G., V. Gallo, E. Quinsal-Keel, and H. MaBouDi. 2025. Bumble bee visual learning: simple solutions for complex stimuli. Animal Behaviour 221:123070.

Hartig, F. 2020. DHARMa: residual diagnostics for hierarchical (multi-level/mixed) regression models. R package version 0.3.3.0.

Hendriksma, H. P., K. L. Oxman, and S. Shafir. 2014. Amino acid and carbohydrate tradeoffs by honey bee nectar foragers and their implications for plant–pollinator interactions. Journal of Insect Physiology 69:56–64.

Herrera, C. M., R. Pérez, and C. Alonso. 2006. Extreme intraplant variation in nectar sugar composition in an insect-pollinated perennial herb. American Journal of Botany 93:575–581.

Hiebert, S. M., and W. A. Calder. 1983. Sodium, potassium, and chloride in floral nectars: energy-free contributions to refractive index and salt balance. Ecology 64:399–402.

Huang, Z.-H., H.-L. Liu, and S.-Q. Huang. 2015. Interspecific pollen transfer between two coflowering species was minimized by bumblebee fidelity and differential pollen placement on the bumblebee body. Journal of Plant Ecology 8:109–115.

Ivushkin, K., H. Bartholomeus, A. K. Bregt, A. Pulatov, B. Kempen, and L. de Sousa. 2019. Global mapping of soil salinity change. Remote Sensing of Environment 231:111260.

Jaiswal, B., S. Singh, S. B. Agrawal, E. Lokupitiya, and M. Agrawal. 2022. Improvements in soil physical, chemical and biological properties at natural saline and non-saline sites under different management practices. Environmental Management 69:1005–1019.

Janssens, S., T. L. P. Couvreur, A. Mertens, G. Dauby, L.-P. Dagallier, S. Vanden Abeele, F. Vandelook, M. Mascarello, H. Beeckman, M. Sosef, V. Droissart, M. Van Der Bank, O. Maurin, W. Hawthorne, C. Marshall, M. Réjou-Méchain, D. Beina, F. Baya, V. Merckx, B. Verstraete, and O. Hardy. 2020. A large-scale species level dated angiosperm phylogeny for evolutionary and ecological analyses. Biodiversity Data Journal 8:e39677.

Kaspari, M. 2020. The seventh macronutrient: how sodium shortfall ramifies through populations, food webs and ecosystems. Ecology Letters 23:1153–1168.

Kaspari, M. 2021. The invisible hand of the periodic table: how micronutrients shape ecology. Annual Review of Ecology, Evolution and Systematics 52:199–219.

Kaspari, M., S. P. Yanoviak, and R. Dudley. 2008. On the biogeography of salt limitation: A study of ant communities. PNAS 105:17848–17851.

Knauer, A. C., and F. P. Schiestl. 2015. Bees use honest floral signals as indicators of reward when visiting flowers. Ecology Letters 18:135–143.

Lau, P. W., and J. C. Nieh. 2016. Salt preferences of honey bee water foragers. Journal of Experimental Biology 219:790–796.

Lenth, R. V., B. Bolker, P. Buerkner, I. Giné-Vázquez, M. Hervé, M. Jung, J. Love, F. Miguez, J. Piaskowski, H. Riebl, and H. Singmann. 2025. emmeans: Estimated Marginal Means, aka Least-Squares Means.

Liu, Y., S. Dunker, W. Durka, C. Dominik, J. M. Heuschele, H. Honchar, P. Hoffmann, M. Musche, R. J. Paxton, J. Settele, and O. Schweiger. 2024. Eco-evolutionary processes shaping floral nectar sugar composition. Scientific Reports 14.

Lovett, D. H., and D. E. Carr. 2024. The potential for elevated soil salinity to enhance the ecological trap effect of roadside pollinator habitat. Journal of Insect Conservation 28:103–111.

Molleman, F. 2010. Puddling: from natural history to understanding how it affects fitness. Entomologia Experimentalis et Applicata 134:107–113.

Morales, C. L., and A. Traveset. 2008. Interspecific pollen transfer: magnitude, prevalence and consequences for plant fitness. Critical Reviews in Plant Sciences 27:221–238.

Nepi, M. 2017. New perspectives in nectar evolution and ecology: simple alimentary reward or a complex multiorganism interaction? Acta Agrobotanica 70.

Nepi, M., D. A. Grasso, and S. Mancuso. 2018. Nectar in plant–insect mutualistic relationships: from food reward to partner manipulation. Frontiers in Plant Science 9.

Nicolson, S. W. 1990. Osmoregulation in a nectar-feeding insect, the carpenter bee *Xylocopa capitata*: water excess and ion conservation. Physiological Entomology 15:433–440.

Nicolson, S. W. 2022. Sweet solutions: nectar chemistry and quality. Philosophical Transactions of the Royal Society B: Biological Sciences 377:20210163.

Nicolson, S. W., M. Nepi, and E. Pacine. 2007. Nectaries and nectar. 1 edition. Springer Dordrecht.

Nicolson, S. W., and R. W. Thornburg. 2007. Nectar chemistry. Pages 215–264 in S. W. Nicolson, M. Nepi, and E. Pacini, editors. Nectaries and Nectar. Springer, Netherlands.

Nicolson, S. W., and P. V. W-Worswick. 1990. Sodium and potassium concentrations in floral nectars in relation to foraging by honey bees. South African Journal of Zoology 25:93–96.

Orme, D., R. P. Freckleton, G. Thomas, T. Petzoldt, S. Fritz, N. Isaac, and W. Pearse. 2018. caper: Comparative Analyses of Phylogenetics and Evolution in R.

Ornelas, J. F., M. Ordano, and C. Lara. 2007. Nectar removal effects on seed production in Moussonia deppeana (Gesneriaceae), a hummingbird-pollinated shrub. Ecoscience 14:117–123.

Ortiz, P. L., P. Fernández-Díaz, D. Pareja, M. Escudero, and M. Arista. 2020. Do visual traits honestly signal floral rewards at community level? Functional Ecology.

Pacini, E., and M. Nepi. 2007. Nectar production and presentation. Pages 167–214 in S. W. Nicolson, M. Nepi, and E. Pacini, editors. Nectaries and nectar. Springer Dordrecht.

Parachnowitsch, A. L., J. S. Manson, and N. Sletvold. 2019. Evolutionary ecology of nectar. Annals of Botany 123:247–261.

Paradis, E., and K. Schliep. 2019. ape 5.0: an environment for modern phylogenetics and evolutionary analyses in R. Bioinformatics 35:526–528.

Pennell, M. W., J. M. Eastman, G. J. Slater, J. W. Brown, J. C. Uyeda, R. G. Fitzjohn, M. E. Alfaro, and L. J. Harmon. 2014. geiger v2.0: an expanded suite of methods for fitting macroevolutionary models to phylogenetic trees. Bioinformatics 30:2216–2218.

Petanidou, T., A. Van Laere, W. N. Ellis, and E. Smets. 2006. What shapes amino acid and sugar composition in Mediterranean floral nectars? Oikos 115:155–169.

POWO. 2024. Plants of the World Online. Facilitated by the Royal Botanic Gardens, Kew. Available from https://powo.science.kew.org.

Prather, C. M., A. N. Laws, J. F. Cuellar, R. W. Reihart, K. M. Gawkins, and S. C. Pennings. 2018. Seeking salt: herbivorous prairie insects can be co-limited by macronutrients and sodium. Ecology Letters 21:1467–1476.

Pyke, G. H. 1991. What does it cost a plant to produce floral nectar? Nature 350:58–59.

Pyke, G. H. 2010. Optimal foraging and plant–pollinator co-evolution. Pages 596-600 in M. D. Breed and J. Moore, editors. Encyclopedia of Animal Behavior. Academic Press, Oxford.

Pyke, G. H. 2016. Floral nectar: pollinator attraction or manipulation? Trends in Ecology & Evolution 31:339–341.

Pyke, G. H., and Z.-X. Ren. 2023. Floral nectar production: what cost to a plant? Biological Reviews 98:2078–2090.

Raubenheimer, D., and S. J. Simpson. 2004. Organismal stoichiometry: quantifying non-independence among food components. Ecology 85:1203–1216.

Raubenheimer, D., and S. J. Simpson. 2018. Nutritional ecology and foraging theory. Current Opinion in Insect Science 27:38–45.

Revell, L. J. 2012. phytools: an R package for phylogenetic comparative biology (and other things). Methods in Ecology and Evolution 3:217–223.

Ruedenauer, F., J. Spaethe, and S. Leonhardt. 2016. Hungry for quality—individual bumblebees forage flexibly to collect high-quality pollen. Behavioral Ecology and Sociobiology 70.

Russell, A. L., S. L. Buchmann, and D. R. Papaj. 2017. How a generalist bee achieves high efficiency of pollen collection on diverse floral resources. Behavioral Ecology 28:991–1003.

Russell, A. L., D. W. Kikuchi, N. W. Giebink, and D. R. Papaj. 2020. Sensory bias and signal detection trade-offs maintain intersexual floral mimicry. Philosophical Transactions of the Royal Society B: Biological Sciences 375:20190469.

Schiestl, F. P., and S. D. Johnson. 2013. Pollinator-mediated evolution of floral signals. Trends in Ecology & Evolution 28:307–315.

Simpson, B. B., and J. L. Neff. 1981. Floral rewards: alternatives to pollen and nectar. Annals of the Missouri Botanical Garden 68:301–322.

Simpson, S. J., and D. Raubenheimer. 1993. A multi-level analysis of feeding behaviour: the geometry of nutritional decisions. Philosophical Transactions: Biological Sciences 342:381–402.

Snell-Rood, E. C., A. Espeset, C. J. Boser, W. A. White, and R. Smykalski. 2014. Anthropogenic changes in sodium affect neural and muscle development in butterflies. PNAS 111:10221–10226.

Southwick, E. E. 1984. Photosynthate allocation to floral nectar: a neglected energy investment. Ecology 65:1775–1779.

Stevenson, P. C. 2020. For antagonists and mutualists: the paradox of insect toxic secondary metabolites in nectar and pollen. Phytochemistry Reviews 19:603–614.

Takimoto, G., K. Kagawa, T. Satow, and T. Sakamoto. 2022. Increased floral rewards due to local adaptation drives plant ecological speciation via learned preferences of pollinators. The American Naturalist 200:834–845.

Team, R. C. 2022. R: A Language and Environment for Statistical Computing. R Foundation for Statistical Computing:https://www.R-project.org/.

Tiedge, K., and G. Lohaus. 2017. Nectar sugars and amino acids in day- and night-flowering Nicotiana species are more strongly shaped by pollinators’ preferences than organic acids and inorganic ions. Plos One 12:e0176865.

Tung Ho, L. s., and C. Ané. 2014. A linear-time algorithm for Gaussian and non-Gaussian trait evolution models. Systematic Biology 63:397–408.

Vannette, R. L. 2020. The floral microbiome: plant, pollinator, and microbial perspectives. Annual Review of Ecology, Evolution, and Systematics 51:363–386.

VanValkenburg, E., T. Gonçalves Souza, N. J. Sanders, and P. CaraDonna. 2024. Sodium-enriched nectar shapes plant–pollinator interactions in a subalpine meadow. 14:e70026.

Varassin, I. G., J. R. Trigo, and M. Sazima. 2001. The role of nectar production, flower pigments and odour in the pollination of four species of Passiflora (Passifloraceae) in south-eastern Brazil. Botanical Journal of the Linnean Society 136:139–152.

Vaudo, A. D., H. M. Patch, D. A. Mortensen, J. F. Tooker, and C. M. Grozinger. 2016. Macronutrient ratios in pollen shape bumble bee (*Bombus impatiens*) foraging strategies and floral preferences. Proceedings of the National Academy of Sciences 113:E4035–E4042.

Welti, E. A. R., and M. Kaspari. 2021. Sodium addition increases leaf herbivory and fungal damage across four grasslands. Functional Ecology 35:1212–1221.

Welti, E. A. R., N. J. Sanders, K. M. De Beurs, and M. Kaspari. 2019. A distributed experiment demonstrates widespread sodium limitation in grassland food webs. Ecology 100:e02600.

Wickham, H. 2016. ggplot2: elegant graphics for data analysis. R package version 3.5.1.

