## Supplementary material for "Salt sweetens the deal: bumble bees prefer sodium-enriched nectar when its sugar concentration is low": Analyses and datasets: readme.md

\# Salty nectar

Authors: Amanda V da Silva¹ \& Avery L. Russell²

Affiliation: Universidade Federal do ABC, ²,




\----------------------------------------------------------------------------------------------

\# General information


\## EXPERIMENT

\### Date of data collection: June 2023

\### Keywords used to describe the data topic: salty nectar, sodium, floral nectar


\## LITERATURE REVIEW

\### Date of data collection: Last updated on May 14th, 2026

\### Keywords used to describe the data topic: salty nectar, sodium, floral nectar


\### Language information: English


\### Information about funding sources that supported the collection of the data: FAPESP BEPE and PhD


\----------------------------------------------------------------------------------------------

\# DATA AND FILE OVERVIEW


STATS\_salt\_concentration\_plants.XLSX


\### Description

&#x09;Sheet1 (data): It contains data on salt concentration found on live flowers of many species of plants. This data was gathered from the literature review.

&#x09;Sheet2 (metadata): It contains explanation on the data and conversions

\### Date(s) that the file(s) was updated (versioned) and the nature of the update(s): May 18th 2026


STATS\_salt\_affect\_nectar\_pref\_07-11-2023-AVS.XLSX


\### Description

&#x09;Sheet1 (data): It contains data on the experiment in which we offered sugar at low concentration (20%) with or without sodium or sugar at high concentration (50%) with or without sodium

&#x09;Sheet2 (metadata): It contains explanation on the data and conversions

\### Date that the file was created: September 26th, 2023


STATS\_bee-head-widths\_7-3-2023\_ALR.XLSX

\### Description

&#x09;Sheet1 (data): It contains measurements of the bee head width used in the experiment

&#x09;Sheet2 (metadata): It contains explanation on the data and conversions

\### Date that the file was created: September 26th, 2023


angiosperm\_phylogram\_janssens2020.tre

\### Description

&#x09;Angiosperm phylogenetic tree by Janssens et al 2020


sodium vs sugar.R

\### Description

&#x09;Analysis to test the impact of salt addition on bees preference

\### Date of creation: October 2025

\### Last update: August, 2026


sodium on floral nectar.R

\### Description

&#x09;Phylogenetic analysis to test whether plants that offer nectar with more sodium reduces the investment in sugar

\### Date of creation: May, 2026


\----------------------------------------------------------------------------------------------

\# SHARING AND ACCESS INFORMATION


\### Links to publications


\----------------------------------------------------------------------------------------------

\# METHODOLOGICAL INFORMATION


\### Description of methods for data collection

\#### Experiment

We divided 36 bumble bees (Bombus impatiens) into two treatment groups, each with two subtreaments. Across all trials, we set up a 5x4 horizontal array of equal numbers of cleaned yellow and blue artificial flowers. Each flower received an artificial nectary into which 3 µL of sterile artificial nectar solution was added. The two treatment groups differed in terms of sucrose (20 or 50% w/v). The subtreaments differed in terms of which of one of the two colors received nectar with 0.1% NaCl w/v.


\#### Literature review

We used the following keywords to search for papers quantifying the amount of sodium and sugar in the nectar of live flowers: (1) salt AND floral nectar; (2) sodium AND floral nectar; (3) NaCl AND floral nectar; (4) (sodium OR salt OR NaCL or ion) AND floral nectar composition; and (5) floral nectar AND (sodium OR salt OR NaCl or ion). We searched on Web of Science on May 14th, 2026. We included all studies that provided data on sodium and sugar content. For each study, we collected plant species, sugar and sodium content.


\### Author contributions: Amanda Vieira da Silva designed the study, collected the data, performed statistical analysis, and wrote the first draft of the manuscript. Avery L Russell designed the study, collected the data, interpreted the data and revised drafts of the manuscript. All authors contributed equally to the final version of the manuscript.


\----------------------------------------------------------------------------------------------

\# DATA INFORMATION


EXCEL SHEETS 


STATS\_salt\_concentration\_plants.XLSX


\### Count of number of variables, and number of cases or rows

Number of variables: 37

Number of observations: 445


\### Variable list, including full names and definitions (spell out abbreviated words) of column headings for tabular data

\* ID: Unique identification for each study	

\* author\_year: Authors and year of the original study	

\* title: Title of the study

\* species: Plant species used in the study	

\* genus: Plant genera used in the study	

\* family: Plant family used in the study	

\* main\_pollinator: Main pollinator recorded

\* condition: Plant cultivated or under field conditions

\* habitat: Which habitat plants were recorded

\* locality: Locality of the study

\* coordinates: Geographic coordinates of the study

\* nectar\_volume\_mean\_ul: Nectar volume of the flower (mean) (uL)

\* nectar\_volume\_se: Standard error of the nectar volume of the flower

\* sample\_size: Sample size of nectar volume

\* sugar: Mean amount of sugar reported

\* sugar\_sd: Standard deviation of sugar

\* sugar\_unity: Measurement unity of sugar

\* sugar\_mM: Only sugar reported in mM

\* method\_sugar: Method to evaluate sugar content

\* sucrose: Amount of sucrose reported

\* sucrose\_sd: Standard deviation of sucrose reported

\* sucrose\_unit: Measurement unity of sucrose

\* glucose: Amount of glucose reported

\* glucose\_sd: Standard deviation of glucose reported

\* glucose\_unit: Measurement unity of glucose

\* fructose: Amount of fructose reported

\* fructose\_sd: Standard deviation of fructose reported

\* fructose\_unit: Measurement unity of fructose

\* sample\_size\_sodium: Sample size of sodium content

\* sodium: Amount of sodium reported

\* sodium\_sd: Standard deviation of sodium reported

\* sodium\_se: Standard error of sodium reported

\* sodium\_95IC: 95% CI of sodium reported

\* sodium\_unity: Measurement unity of sodium

\* sodium\_mM: Converted sodium values to mM

\* sodium\_mass\_nectar\_volume: Amount of sodium available on floral nectar

\* Obs: Observations


\### Definitions for codes used to record missing data

NA or empty cell


STATS\_salt\_affect\_nectar\_pref\_07-11-2023-AVS.XLSX


\### Count of number of variables, and number of cases or rows

Number of variables: 14

Number of observations: 1522


\### Variable list, including full names and definitions (spell out abbreviated words) of column headings for tabular data

\* observers: Researchers who made the observation

\* date: date of the trial (DD/MM/YYYY)

\* begin\_time: hour in which the trial began (24h)

\* end\_time: hour in which the trial ended (24h)

\* colony: ID of the colony

\* bee\_ID: ID of the bee

\* sugar\_concentration: percentage of sugar used in that trial

\* salt\_color: color of the treatment that represents addition of salt (1%)

\* treatment: combination of sugar concentration and salt

\* n\_visits: number of visits performed by a bee

\* flower: color of the flower visited by the bee 

\* flower\_salt: was the visit performed on a flower with salt addition? (yes or no)

\* behavior: drink (bees drank the nectar solution), taste (bees inserted the tongue into the feeder, but did not drink), land (land on a flower), empty (there was no nectar on the feeder), return (bee returned and visited the previous visited flower), return empty (return to the flower and it was empty)

\* comments: additional comments


\### Definitions for codes used to record missing data

NA


STATS\_bee-head-widths\_7-3-2023\_ALR


\### Count of number of variables, and number of cases or rows

Number of variables: 7

Number of observations: 41


\### Variable list, including full names and definitions (spell out abbreviated words) of column headings for tabular data

\* bee\_ID: ID of the bee

\* treatment: percentage of sugar used in that trial

\* subtreatment: color of the treatment that represents addition of salt (1%)

\* file\_name: name of the file picture saved

\* head\_width\_px: width of bee head in pixels

\* head\_width\_mm: width of bee head in mm

\* 1\_mm\_px: conversion of 1 mm to pixels


R SCRIPTS


\* sodium on floral nectar.R: phylogenetic comparative analysis of the relationship between sugar and sodium concentration in nectar of live flowers


\* sodium vs sugar.R: analysis on the effect of sodium-addition and sugar concentration on bee preference
