## Supplementary material for "Salt sweetens the deal: bumble bees prefer sodium-enriched nectar when its sugar concentration is low": Analyses and datasets: sodium on floral nectar.R

```
## Relationship between salt and sugar concentration ##
## May 18th, 2026 ## 
## By: Amanda Vieira da Silva ##
## ##
## GitHub: @mandyviesi ##

# PACKAGES ---------------------------

# Data manipulation
library(dplyr)

# Phylogenetic analysis
library(V.PhyloMaker2)
library(phytools)
library(ape)
library(phylolm)

#Figures
library(ggplot2)

# DATA ----------------------------------------

## Nectar traits-------------------------------

# Importing data
nectar <- readxl::read_xlsx("STATS_salt_concentration_plants.xlsx", na = "NA")

# String as factors
nectar <- nectar |> 
  mutate_if(is.character, as.factor)

# Checking it
glimpse(nectar)
head(nectar)
summary(nectar)


## Tree -------------------------------------------------

# Angiosperm phylogenetic tree
# Based on: Janssens et al 2020
angiotree <- read.nexus("angiosperm_phylogram_janssens2020.tre")

angiotree

# DESCRIPTIVE STATS ------------------------

# Number of studies: 15
nectar |> 
  distinct(ID_study) |> 
  count()

# Number of studies that provided that on both sodium AND sugar content on floral nectar
nectar |> 
  filter(!is.na(sugar), !is.na(sodium)) |> 
  distinct(ID_study) |> 
  count()

# Calculating mean sodium and sugar concentration on studies regardless of the fact they had data for both traits

# Sodium
nectar |> 
  summarise(meanNa = mean(sodium_mM, na.rm = TRUE),
            medianNa = median(sodium_mM, na.rm = TRUE),
            SDNa = sd(sodium_mM, na.rm = TRUE),
            minNa = min(sodium_mM, na.rm = TRUE),
            maxNa = max(sodium_mM, na.rm = TRUE))

# Sugar
nectar |> 
  summarise(meansugar = mean(sugar_mM, na.rm = TRUE),
mediansugar = median(sugar_mM, na.rm = TRUE),
SDsugar = sd(sugar_mM, na.rm = TRUE),
minsugar = min(sugar_mM, na.rm = TRUE),
maxsugar = max(sugar_mM, na.rm = TRUE))

# Creating a dataframe from studies that provided that on both sodium AND sugar content on floral nectar
nectar_stats <- nectar |> 
  filter(!is.na(sugar), !is.na(sodium))

# Number of plant species for which we have data on both sodium and sugar content on floral nectar
nectar_stats |> 
  group_by(species) |> 
  count() |> 
  arrange(desc(n))

# Number of plant genera for which we have data on both sodium and sugar content on floral nectar 
nectar_stats |>   
  group_by(genus) |> 
  count() |> 
  arrange(desc(n))

# # Number of plant genera for which we have data on both sodium and sugar content on floral nectar
nectar_stats |> 
  group_by(family) |> 
  count() |> 
  arrange(desc(n))

# Mean and SD sodium concentration
nectar_stats |> 
  summarise(meanNa = mean(sodium_mM, na.rm = TRUE),
            SDNa = sd(sodium_mM, na.rm = TRUE),
            medianNa = median(sodium_mM, na.rm = TRUE),
            minNa = min(sodium_mM, na.rm = TRUE),
            maxNa = max(sodium_mM, na.rm = TRUE))

# Mean and SD sugar concentration
nectar_stats |> 
  summarise(meanSugar = mean(sugar_mM, na.rm = TRUE),
            SDSugar = sd(sugar_mM, na.rm = TRUE),
            medianSugar = median(sugar_mM, na.rm = TRUE),
            minSugar = min(sugar_mM, na.rm = TRUE),
            maxSugar = max(sugar_mM, na.rm = TRUE))

# Accounting for nectar volume
nectar |> 
  summarise(meanNaNectar = mean(sodium_mass_nectar_volume, na.rm = TRUE),
            sdNaNectar = sd(sodium_mass_nectar_volume, na.rm = TRUE))


# Analysis  -------------------------------

## Species level --------------------------
# Creating a dataframe with one value per species
nectarsp <- nectar |> 
  filter(!is.na(sugar_mM), !is.na(sodium_mM)) |> 
  group_by(species) |> 
  summarise(sugar = mean(sugar_mM),
            sodium = mean(sodium_mM))

# Row names
rownames(nectarsp) <- stringr::str_to_sentence(nectarsp$species)

# Pruning the tree to the data
check <- geiger::name.check(phy = angiotree, data = nectarsp)

# Drop tips from the tree
nectartree <- drop.tip(angiotree, check$tree_not_data)

# Subseting data
nectarspdata <- nectarsp[nectartree$tip.label,]

# As data frame
nectarspdata <- as.data.frame(nectarspdata)

# Rownames as written in the tree
rownames(nectarspdata) <- stringr::str_to_sentence(nectarspdata$species)

# Reorganizing the dataframe
nectarspdata <- nectarspdata[match(nectartree$tip.label, rownames(nectarspdata)), ]

rownames(nectarspdata) == nectartree$tip.label

## Models --------------------------------------

# BM model
phylobm <- phylolm(formula = sodium ~ sugar, data = nectarspdata, phy = nectartree, model = "BM")
summary(phylobm)
plot(phylobm)

# Creating dataframe for figure
df_fig <- data.frame(sugar = seq(min(nectarspdata$sugar),
                                      max(nectarspdata$sugar),
                                      length.out = nrow(nectarspdata)))

# Adding the predicted values
df_fig$pred <- predict(phylobm, newdata = df_fig)

# Figure - no correlation: dashed line
tiff("fig4-may2026_model.tif", width = 1500, height = 1500, res = 300, compression = "lzw")
ggplot(nectarspdata, aes(x = sugar, y = sodium)) +
  geom_point(size = 4, pch = 20, alpha = 0.5, color = "gray") +
  geom_line(data = df_fig, aes(y = pred), color = "#E35205", linewidth = 1, linetype = "dashed") +
  labs(x = "Mean sugar (mM)", y = "Mean sodium (mM)") +
  theme(axis.text = element_text(size = 14, color = "black"),
        axis.title = element_text(size = 16),
        axis.line = element_line(color = "black"),
        panel.background = element_rect(fill = "white"),
        aspect.ratio = 1)
dev.off()

# Figure using real data
# Nectar and sugar mean per species
nsmean <- nectar2 |> 
  group_by(species) |> 
  summarise(sugar_mM = mean(sugar_mM, na.rm = TRUE),
            sodium_mM = mean(sodium_mM, na.rm = TRUE))

nsmean <- na.omit(nsmean)

tiff("fig4-may2026_realdata.tif", width = 1500, height = 1500, res = 300, compression = "lzw")
ggplot(nsmean, aes(x = sugar_mM, y = sodium_mM)) +
  geom_point(size = 4, pch = 20, alpha = 0.5, color = "gray") +
  geom_line(data = df_fig, aes(y = pred, x = sugar), color = "#E35205", linewidth = 1, linetype = "dashed") +
  labs(x = "Mean sugar (mM)", y = "Mean sodium (mM)") +
  scale_y_continuous(limits = c(0,8)) +
  theme(axis.text = element_text(size = 14, color = "black"),
        axis.title = element_text(size = 16),
        axis.line = element_line(color = "black"),
        panel.background = element_rect(fill = "white"),
        aspect.ratio = 1)
dev.off()
```
