## Supplementary material for "Salt sweetens the deal: bumble bees prefer sodium-enriched nectar when its sugar concentration is low": Analyses and datasets: sodium vs sugar.R

```
## Effect of sodium-addition and sugar concentration on bees preference ##
## August, 2026 ## 
## By: Amanda Vieira da Silva ##
## ##
## GitHub: @mandyviesi ##

# Packages ---------------------------

# Data manipulation
library(dplyr)

# Models
library(glmmTMB)
library(emmeans)
library(car)
library(DHARMa)
library(sjPlot)

#Figures
library(ggplot2)

# Data -------------------------------

## Preference essay data -------------

# Data
pref <- readxl::read_xlsx("STATS_salt_affect_nectar_pref_07-11-2023-AVS.xlsx")

# Checking it out
glimpse(pref)

# Changing begin_time and end_time to time
pref <- pref |> 
  mutate(begin_time = lubridate::hm(begin_time),
         end_time = lubridate::hm(end_time))


# Changing character variables to factor
pref <- pref |> 
  mutate(across(where(is.character), factor))

# Checking it out: ok
glimpse(pref)

## Bees width head -----------------------

# Data
width <- readxl::read_xlsx("STATS_bee-head-widths_7-3-2023_ALR.xlsx", sheet = 2, na = "NA")

# Checking it out
glimpse(width)

# Changing character as factor
width <- width |> 
  mutate(across(where(is.character), factor))

# Checking it out
glimpse(width)

# Selecting variables 
bee_size <- width |> 
  select(bee_ID, head_width_mm)

# Lower case
bee_size$bee_ID <- tolower(bee_size$bee_ID)

## Joining the data -----------------------

# Joining
salty <- pref |> 
  left_join(bee_size, by = "bee_ID")

# Checking it out
glimpse(salty)

# Removing 10% sugar treatment
salty <- salty |> 
  filter(sugar_concentration != 0.1)

# Removing datasets we will not use anymore
rm(bee_size)
rm(pref)
rm(width)

# Descriptive stats ------------------------------

# Checking dataset
glimpse(salty)

# Number of bees
salty |> 
  distinct(bee_ID) |> 
  count()

# Number of bees per treatment
salty |> 
  group_by(treatment) |> 
  distinct(bee_ID) |> 
  count()

# Mean number of visits per bee
# Drink as a number
salty <- salty |> 
  mutate(drink2 = case_when(
    behavior == "drink" ~ 1,
    behavior == "return" ~ 1,
    behavior != "drink" | behavior != "return" ~ 0))

# Calculating how much time each bee visit and drink nectar 
drinking_bee <- salty |> 
  group_by(bee_ID) |> 
  summarise(sum_bee = sum(drink2))

# Calculating mean and standard deviation
drinking_bee |> 
  summarise(mean = mean(sum_bee),
            sd = sd(sum_bee))


# Q1: Proportion of landing ------------------------

# Hypothesis: bees will be more prone to land and reject flowers with low-quality rewards and no salt when compared to flowers with low-quality rewards and salt. Also, bees will be equally prone to land and reject flowers with high-quality rewards, regardless of salt.
# Adding landing as a variable in the dataset 
# Code: 
## Land but did not drink (land followed by rejection): 1  whereas landing and accepting (considering that bees need to land before drinking): 0 
unique(salty$behavior)

for(i in 1:nrow(salty)){
  if(salty$behavior[i] == "land" | salty$behavior[i] == "return land"){ 
    salty$land[i] <- 1
  }else{
    salty$land[i] <- 0
  }
}

# Almost 4x more drinking than landing and rejecting
table(salty$land)

# Model 1: Binomial error distribution
# Checking whether flower salt and sugar concentration are factors
class(salty$flower_salt)
class(salty$sugar_concentration)

# Transforming sugar concentration
salty$sugar_concentration <- as.factor(salty$sugar_concentration)

#Model
mod1_land_binomial <-  glmmTMB(land ~ flower_salt * sugar_concentration + (1|bee_ID), 
                               family=binomial, 
                               data=salty)

# Model summary
summary(mod1_land_binomial) 

# Checking singularity
performance::check_singularity(mod1_land_binomial)

# Checking convergence
performance::check_convergence(mod1_land_binomial)

# Checking homocedasticity
mod1_simulation <-  simulateResiduals(fittedModel = mod1_land_binomial, plot = T)

# Checking overdispersion
testDispersion(mod1_land_binomial)

# Checking zero inflation
testZeroInflation(mod1_land_binomial)

# Generating p-values
Anova(mod1_land_binomial, type = "2")

# Posthoc comparisons
emmeans(mod1_land_binomial, 
        pairwise~flower_salt | sugar_concentration,
        adjust="tukey")

# Effect size calculation
emm1 <- emmeans(mod1_land_binomial, ~ flower_salt | sugar_concentration, type = "response")
emm1
eff_size(emm1, sigma = sigma(mod1_land_binomial ), edf = df.residual( mod1_land_binomial))

# Calculating proportion for figures
land <- salty |> 
  select(bee_ID, flower_salt, sugar_concentration, land) |> 
  group_by(flower_salt, sugar_concentration) |> 
  summarise(
    total = n(),
    land = sum(land, na.rm = TRUE),
    proportion = land/total
  )

# Calculating standard error of the proportion
land
land$sep <- sqrt(land$proportion*(1-land$proportion)/36)

# Calculating effect sizes
land

## Low sucrose concentrations
land$proportion[1] / land$proportion[3]

# Figure
fig2a <- ggplot(land, aes(y = proportion, x = as.factor(sugar_concentration), color = flower_salt)) +
  geom_point(size = 3, position = position_dodge(width = 0.3)) +
  geom_errorbar(aes(ymin = proportion - sep, ymax = proportion + sep), width = 0.2, position = position_dodge(width = 0.3)) +
  scale_color_manual(name = "Salt addition", 
                     values = c("#0093B2", "#E35205"),
                     labels = c("No", "Yes")) +
  scale_y_continuous(name = "Proportion of landing") +
  scale_x_discrete(name = "Sucrose concentration", 
                   labels = c("20%", "50%")) +
  annotate(geom = "text", label = "*", x = 1, y = 0.35, size = 5) +
  theme(panel.background = element_blank(),
        axis.line = element_line(color = "black"),
        axis.text = element_text(color = "black", size = 12),
        axis.title = element_text(color = "black", size = 14),
        legend.position = "top",
        aspect.ratio = 1)

# Q2: Proportion of tasting ---------------------

# Hypothesis: bees will be more prone to reject low-quality nectar without salt than high-quality nectar, regardless of salt in the high-quality nectar.

# Adding taste as a variable in the dataset 
# Code: 
## Taste but did not drink (taste followed by rejection): 1  whereas taste and accepting (considering that bees need to taste before drinking): 0
unique(salty$behavior)
salty <- salty |> 
  mutate(taste = case_when(
    behavior == "taste" ~ 1,
    behavior == "drink" ~ 0,
    behavior == "return" ~ 0))

# Creating a dataframe
taste <- salty |> 
  select(taste, flower_salt, sugar_concentration, bee_ID, n_visits)

# Removing NA
taste <- na.omit(taste)

# Almost 9x more tasting than drinking
table(taste$taste)

# Model 3: Binomial error distribution
class(taste$flower_salt)
class(taste$sugar_concentration)

mod2_taste_binomial <-  glmmTMB(taste ~ flower_salt* sugar_concentration + (1|bee_ID), 
                                family=binomial, 
                                data=taste)

# Model summmary
summary(mod2_taste_binomial) 

# Checking singularity
performance::check_singularity(mod2_taste_binomial)

# Checking convergence
performance::check_convergence(mod2_taste_binomial)

# Checking homocedasticity
mod2_simulation <-  simulateResiduals(fittedModel = mod2_taste_binomial, plot = T)

# Checking dispersion
testDispersion(mod2_taste_binomial)

# Checking zero inflation
testZeroInflation(mod2_taste_binomial)

# Generating p-values
Anova(mod2_taste_binomial, type="2")

# Post-hoc comparisons
emmeans(mod2_taste_binomial, 
        pairwise~flower_salt | sugar_concentration,
        adjust="tukey")

# Effect size calculation
emm2 <- emmeans(mod2_taste_binomial, ~ flower_salt | sugar_concentration, type = "response")
emm2
eff_size(emm2, sigma = sigma(mod2_taste_binomial), edf = df.residual( mod2_taste_binomial))

# Calculating proportion for figures
taste_fig <- taste |> 
  group_by(flower_salt, sugar_concentration) |> 
  summarise(
    total = n(),
    taste = sum(taste, na.rm = TRUE),
    proportion = taste/total
  )
View(taste)

# Calculating standard error of the proportion
taste_fig
taste_fig$sep <- sqrt(taste_fig$proportion*(1-taste_fig$proportion)/36)

# Calculating effect sizes
taste_fig$proportion[1] / taste_fig$proportion[3]

# Figure
fig2b <- ggplot(taste_fig, aes(y = proportion, x = as.factor(sugar_concentration), col = flower_salt)) +
  geom_point(size = 3,  position = position_dodge(width = 0.3)) +
  geom_errorbar(aes(ymin = proportion - sep, ymax = proportion + sep), width = 0.2,  position = position_dodge(width = 0.3)) +
  scale_color_manual(name = "Salt addition", 
                     values = c("#0093B2", "#E35205"),
                     labels = c("No", "Yes")) +
  scale_y_continuous(name = "Proportion of tasting") +
  scale_x_discrete(name = "Sucrose concentration", 
                   labels = c("20%", "50%")) +
  annotate(geom = "text", label = "*", x = 1, y = 0.46, size = 5) +
  theme(panel.background = element_blank(),
        axis.line = element_line(color = "black"),
        axis.text = element_text(color = "black", size = 12),
        axis.title = element_text(color = "black", size = 14),
        legend.position = "top",
        aspect.ratio = 1)


# Q3: Proportion of drinking ----------------------------

# Hypothesis: bees will prefer salty nectar and this preference  will be stronger when bees are offered low-quality nectar reward.

# Adding drinking as a variable in the dataset (Code: bees drank: 1, if bees did not drink: 0)
salty <- salty |> 
  mutate(drink = case_when(
    behavior == "drink" ~ 1,
    behavior == "return" ~ 1,
    behavior != "drink" | behavior != "return" ~ 0))

# Model 1: Binomial error distribution
class(salty$flower_salt)
class(salty$sugar_concentration)

mod3_drinking_binomial <-  glmmTMB(drink2 ~ flower_salt*sugar_concentration + (1|bee_ID), 
                                   family=binomial, 
                                   data=salty)

# Model summary
summary(mod3_drinking_binomial) 

# Checking singularity
performance::check_singularity(mod3_drinking_binomial)

# Checking convergence
performance::check_convergence(mod3_drinking_binomial)

# Checking homocedasticity
mod3_simulation <-  simulateResiduals(fittedModel = mod3_drinking_binomial, plot = T)

# Checking dispersion
testDispersion(mod3_drinking_binomial)

# Checking zero inflation
testZeroInflation(mod3_drinking_binomial)

# Generating p-values
Anova(mod3_drinking_binomial, type="2")

# Post-hoc comparisons
emmeans(mod3_drinking_binomial, 
        pairwise~flower_salt | sugar_concentration,
        adjust="tukey")

# Effect size calculation
emm3 <- emmeans(mod3_drinking_binomial, ~ flower_salt | sugar_concentration, type = "response")
emm3
eff_size(emm3, sigma = sigma(mod3_drinking_binomial), edf = df.residual( mod3_drinking_binomial))

# Calculating proportion for figures
drink <- salty |> 
  group_by(flower_salt, sugar_concentration) |> 
  summarise(
    total = n(),
    drink = sum(drink, na.rm = TRUE),
    proportion = drink/total
  )

# Calculating standard error of the proportion
drink
drink$sep <- sqrt(drink$proportion*(1-drink$proportion)/36)

# Calculating effect sizes
drink$proportion[3] / drink$proportion[1]

# Figure
fig2c <- ggplot(drink, aes(y = proportion, x = as.factor(sugar_concentration), col = flower_salt)) +
  geom_point(size = 3, position = position_dodge(width = 0.3)) +
  geom_errorbar(aes(ymin = proportion - sep, ymax = proportion + sep), width = 0.2, position = position_dodge(width = 0.3)) +
  scale_color_manual(name = "Salt addition", 
                     values = c("#0093B2", "#E35205"),
                     labels = c("No", "Yes")) +
  scale_y_continuous(name = "Proportion of drinking") +
  scale_x_discrete(name = "Sucrose concentration", 
                   labels = c("20%", "50%")) +
  annotate(geom = "text", label = "*", x = 1, y = 0.9, size = 5) +
  theme(panel.background = element_blank(),
        axis.line = element_line(color = "black"),
        axis.text = element_text(color = "black", size = 12),
        axis.title = element_text(color = "black", size = 14),
        legend.position = "top",
        aspect.ratio = 1)

# Figure 2 -----------------

# Creating figure 2
tiff("fig2.tif", res = 300, h = 1150, w = 2500)
ggpubr::ggarrange(fig2a, fig2b, fig2c, nrow = 1,
                  common.legend = TRUE,
                  labels = c("(A)", "(B)", "(C)"))
dev.off()

# Q4: Switching to a different flower ---------------

# Hypothesis: bees will be less prone to change to a different flower after tasting flowers with salt, and this change will be stronger when bees are offered low-quality nectar reward:
# GLMM: treatment * sugar as the predictor variable, frequency of changing as the response variable, and bee ID as a random effect (error distribution: Poisson or binomial negative)
glimpse(salty)

# Creating a variable with change
# Code: if bee forage on a flower with a different color from the previous one: switch; if bee forage on a flower with the same color of the previous one: no_switch
salty <- salty |> 
  group_by(bee_ID) |> 
  mutate(switch = ifelse(flower != lag(flower), "switch", "no_switch")) 

# Removing first flower visited (NA)
switch <- salty |> 
  select(switch, flower_salt, sugar_concentration, bee_ID, n_visits, behavior)

switch <- na.omit(switch)

# Subsetting behaviors (only bees who drink and taste)
# Return = drink nectar from the previous visited flower
unique(switch$behavior)

switch <- switch |> 
  filter(behavior == "drink" |
           behavior == "return" |
           behavior == "taste")

# Subsetting data to get bee visit after first switch
switch2 <- switch |> 
  group_by(bee_ID) |> 
  filter(which(switch == "switch")[1] <= row_number())
switch2

# Changing switch variable to numeric
switch2 <- switch2 |> 
  mutate(switch = case_when(
    switch == "no_switch" ~ 0,
    switch == "switch" ~ 1))


# Almost 1.6 times more no switching than switching
table(switch2$switch)

# Model 4: Binomial error distribution
class(switch2$flower_salt)
class(switch2$sugar_concentration)

mod4_switch_binomial <-  glmmTMB(switch ~ flower_salt * sugar_concentration + (1|bee_ID), 
                                 family=binomial, 
                                 data=switch2)

# Checking model assumptions
summary(mod4_switch_binomial) 

# Checking singularity
performance::check_singularity(mod4_switch_binomial)

# Checking convergence
performance::check_convergence(mod4_switch_binomial)

# Checking homocedasticity
mod4_simulation <-  simulateResiduals(fittedModel = mod4_switch_binomial, plot = T)

# Checking dispersion
testDispersion(mod4_switch_binomial)

# Checking zero inflation
testZeroInflation(mod4_switch_binomial)

# Generating p-values
Anova(mod4_switch_binomial, type="2")

# Post-hoc comparisons
emmeans(mod4_switch_binomial, 
        pairwise~flower_salt | sugar_concentration,
        adjust="tukey")

# Effect size calculation
emm4 <- emmeans(mod4_switch_binomial, ~ flower_salt | sugar_concentration, type = "response")
emm4
eff_size(emm4, sigma = sigma(mod4_switch_binomial), edf = df.residual( mod4_switch_binomial))

# Calculating proportion for figures
freqswitch <- switch2 |> 
  group_by(flower_salt, sugar_concentration) |> 
  summarise(
    total = n(),
    switch = sum(switch, na.rm = TRUE),
    proportion = switch/total
  )

# Calculating standard error of the proportion
freqswitch
freqswitch$sep <- sqrt(freqswitch$proportion*(1-freqswitch$proportion)/36)

# Calculating effect sizes
freqswitch$proportion[1] / freqswitch$proportion[3]

# Figure
tiff("fig3.tif", res = 300, h = 1500, w = 1500)
ggplot(freqswitch, aes(y = proportion, x = as.factor(sugar_concentration), col = flower_salt)) +
  geom_point(size = 3, position = position_dodge(width = 0.3)) +
  geom_errorbar(aes(ymin = proportion - sep, ymax = proportion + sep), width = 0.2, position = position_dodge(width = 0.3)) +
  scale_color_manual(name = "Salt addition", 
                     values = c("#0093B2", "#E35205"),
                     labels = c("No", "Yes")) +
  scale_y_continuous(name = "Proportion of switching") +
  scale_x_discrete(name = "Sucrose concentration", 
                   labels = c("20%", "50%")) +
  theme(panel.background = element_blank(),
        axis.line = element_line(color = "black"),
        axis.text = element_text(color = "black", size = 12),
        axis.title = element_text(color = "black", size = 14),
        legend.position = "top",
        aspect.ratio = 1)
dev.off()
```
