## Supplementary material for "Salt sweetens the deal: bumble bees prefer sodium-enriched nectar when its sugar concentration is low": Tables S1

**Table S1.** Estimates of generalized linear mixed models (GLMM) describing bumble bee (*Bombus impatiens*) behavior on artificial flowers with or without sodium addition. ‘Landing’ was when the bee visited the flower (touched the flower with at least 3 of its legs simultaneously*) but* did not extend its proboscis into the nectary; ‘tasting’, when the bee visited and extended its proboscis into the nectary only briefly (<<1 second); ‘drinking’, when the bee visited and extended its proboscis into the nectary for more than 1 second; and ‘switching’ was when the bee switched to a different flower type (one with versus without NaCl in its nectar) by bumble bees.

| **Model** | **Estimate** | **SE** | **Z** | **p-value** |
| --- | --- | --- | --- | --- |
| *Proportion of landing* | | | | |
| Intercept  Salt presence  Sugar concentration  Salt presence: Sugar concentration | -1.084  -0.876  -0.713  0.587 | 0.184  0.233  0.255  0.329 | -5.895  -3.760  -2.793  -1.784 | <0.001  <0.001  <0.001  0.074 |
| *Proportion of tasting* | | | | |
| Intercept  Salt presence  Sugar concentration  Salt presence: Sugar concentration | -0.767  -2.323  -3.220  2.591 | 0.307  0.321  0.531  0.609 | -2.501  -7.235  -6.066  4.259 | 0.012  <0.001  <0.001  <0.001 |
| *Proportion of drinking* | | | | |
| Intercept  Salt presence  Sugar concentration  Salt presence: Sugar concentration | -0.153  1.574  1.542  -1.315 | 0.185  0.205  0.257  0.295 | -0.774  7.689  5.994  -4.450 | 0.439  <0.001  <0.001  <0.001 |
| *Proportion of switching* | | | | |
| Intercept  Salt presence  Sugar concentration  Salt presence: Sugar concentration | -0.123  -0.340  -0.547  0.657 | 0.245  0.220  0.327  0.293 | -0.500  -1.542  -1.673  2.202 | 0.617  0.123  0.094  0.028 |
